# Classifying the activity states of small vertebrates using automated VHF telemetry

**DOI:** 10.1101/2022.03.22.485147

**Authors:** Jannis Gottwald, Raphaël Royauté, Marcel Becker, Tobias Geitz, Jonas Höchst, Patrick Lampe, Lea Leister, Kim Lindner, Julia Maier, Sascha Rösner, Dana G. Schabo, Bernd Freisleben, Roland Brandl, Thomas Müller, Nina Farwig, Thomas Nauss

## Abstract

- The most basic behavioural states of animals can be described as active or passive. However, while high-resolution observations of activity patterns can provide insights into the ecology of animal species, few methods are able to measure the activity of individuals of small taxa in their natural environment. We present a novel approach in which the automated VHF radio-tracking of small vertebrates fitted with lightweight transmitters (< 0.2 g) is used to distinguish between active and passive behavioural states.
- A dataset containing > 3 million VHF signals was used to train and test a random forest model in the assignment of either active or passive behaviour to individuals from two forest-dwelling bat species (*Myotis bechsteinii* (n = 50) and *Nyctalus leisleri* (n = 20)). The applicability of the model to other taxonomic groups was demonstrated by recording and classifying the behaviour of a tagged bird and by simulating the effect of different types of vertebrate activity with the help of humans carrying transmitters. The random forest model successfully classified the activity states of bats as well as those of birds and humans, although the latter were not included in model training (F-score 0.96–0.98).
- The utility of the model in tackling ecologically relevant questions was demonstrated in a study of the differences in the daily activity patterns of the two bat species. The analysis showed a pronounced bimodal activity distribution of *N. leisleri* over the course of the night while the night-time activity of *M. bechsteinii* was relatively constant. These results show that significant differences in the timing of species activity according to ecological preferences or seasonality can be distinguished using our method.
- Our approach enables the assignment of VHF signal patterns to fundamental behavioural states with high precision and is applicable to different terrestrial and flying vertebrates. To encourage the broader use of our radio-tracking method, we provide the trained random forest models together with an R-package that includes all necessary data-processing functionalities. In combination with state-of-the-art open-source automated radio-tracking, this toolset can be used by the scientific community to investigate the activity patterns of small vertebrates with high temporal resolution, even in dense vegetation.

## Introduction

The behaviour of an animal can be fundamentally divided into active and passive (Halle & Stenseth, 2000), with the former requiring a much higher energy expenditure (Rowcliffe et al., 2014). Quantifying the distribution of activity periods throughout the day provides important insights into species’ responses to their environment, foraging strategies, bioenergetics and adaptations (Aschoff, 1966; Torney et al., 2021). Moreover, knowledge of the partitioning of sympatric species along the temporal niche axis can yield insights into the mechanisms that facilitate stable coexistence (Nakabayashi et al., 2021).

Detailed analyses of the activity patterns of individuals requires high-resolution observations (Nathan et al., 2022), which are often difficult to obtain (Williams et al., 2014). The observer’s presence may influence animal behaviour and thus limit conclusions (Crofoot et al., 2010; Isbell & Young, 1993) and continuous observation of elusive or highly mobile species in habitats with dense vegetation is close to impossible (Maffei et al., 2005). While information on medium-sized to large species can be obtained using camera traps (Hughey et al., 2018), GPS transmitters and accelerometers (Kays et al., 2015), as demonstrated in investigations of dynamic habitat and resource use (Wyckoff et al., 2018), behaviour (Freeman et al., 2010) and migration and dispersal (Walton et al., 2018), these devices are of limited use for small (<100 g) animals, due to low detection probabilities, the trade-off between transmitter size and weight, battery life and data-collection intensity (Hallworth & Marra, 2015; Hammond et al., 2016; Wikelski et al., 2007). Newer technical solutions such as the ATLAS system (Nathan et al., 2022) or WBN (Ripperger et al., 2020) allow the tracking of small animals with high temporal and spatial resolution, but the required installation effort and the costs are high.

Very high frequency (VHF) telemetry has been employed in wildlife tracking since the 1960s (Cochran et al., 1965; Lord et al., 1962), with the ongoing miniaturisation of VHF transmitters (< 0.2 g) allowing the tracking of small taxa (body mass < 5 g), ranging from large insects to small vertebrates (Fisher et al., 2020; Naef-Daenzer et al., 2005). Such studies take advantage of the fact that even small movements of tagged animals result in discernible variations in the strength of the received signal (Cochran & Lord, 1963; Kjos & Cochran, 1970) that reflect changes in the angle and distance between the transmitter and receiver (Fig.1). However, collecting reasonable amounts of data on activity bouts using manual radio-telemetry requires an enormous amount of fieldwork (Kjos & Cochran, 1970), which implies a high level of wildlife disturbance (Kenward, 2000; Mech & Barber, 2002) and the risk of missing critical events in the life of the tagged individuals is high (Lambert et al., 2009).

**Fig. 1.**
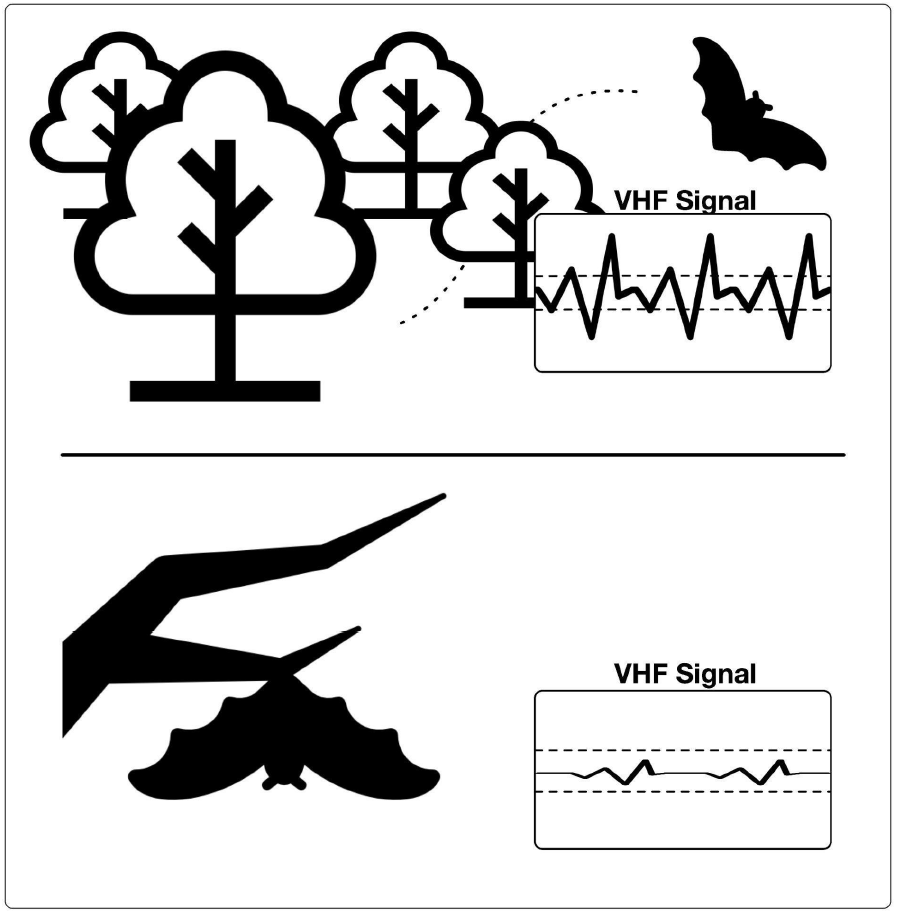
Principle of activity-recognition-based very high frequency (VHF) signal patterns. Top: flying bat; bottom: resting bat. The amplitude and variation of the signal strength over time increase when the tagged individual is moving.

Kays et al. (2011) proposed a method for automatically classifying active and passive behaviour based on a threshold difference in the signal strength of successive VHF signals recorded by a customised automatic radio-tracking system. However, a limitation of the used radio-tracking system is that it can only track one tag at a time, resulting in a low temporal resolution (i.e. a few seconds of observations every 10 min due to switching through frequency channels).

High-throughput tracking systems (<10-s data interval, many individuals at a time) enable ground-breaking research in animal behaviour, evolution and ecology (Nathan et al., 2022). In recent years, with the ongoing development of low-cost open-source solutions, automatic VHF radio-tracking has become broadly available to researchers. These systems allow the tracking of many individuals simultaneously and with a very high temporal resolution (seconds) over the complete tagging period (Gottwald et al., 2019; Höchst et al., 2021; Taylor et al., 2017). Continuous, high-resolution recording of the VHF signals makes the entire signal pattern available for subsequent data analysis.

Machine learning (ML) algorithms are optimised for the recognition of complex patterns in a dataset and are typically robust against factors that influence signal propagation, such as changes in temperature and humidity, physical contact with conspecifics and/or multipath signal propagation (Alade, 2013). Accordingly, a ML model trained with a dataset encompassing the possible diversity of signal patterns related to active and passive behaviour can be expected to perform at least as well as a threshold-based approach.

In this work, we built on the methodology of Kays et al. (2011) by calibrating a ML method,i.e. a random forest model, based on millions of data points representing the behaviours of multiple tagged individuals of two temperate bat species (*Myotis bechsteinii, Nyctalus leisleri*). This strategy was used in conjunction with recent developments in automated radio-telemetry (Gottwald et al., 2019, 2021; Höchst et al., 2021) to develop a toolset that allows researchers to record the activity patterns of even very small species (body mass < 5 g) in their natural habitat and with high resolution. The method was tested by applying it to independent data from bats, humans and a bird species and then comparing the results with those obtained using the threshold-based approach of Kays et al. (2011).

In our method, activity states are recognised with high temporal resolution (< 10 s) and high accuracy. In the following, we provide detailed information on the application of the random forest model and its validation using data on the behaviour of tagged bat and bird individuals generated with an open-source multi-sensor tool (Gottwald & Lampe et al. 2020). In a case study, we demonstrate the use of the approach to detect differences in activity patterns (here, between those of *M. bechsteinii* and *N. leisleri*) and to compare breeding vs. non-breeding status. Detailed information on data processing and analysis is provided, along with an R-package, example scripts and data, all stored in an open data repository.

The scientific contributions of this study are:

1. the development of a random forest model to classify VHF signals obtained from two bat species using an automatic radio-tracking system (tRackIT) into active and passive states, with the validity of the model tested in a bird species and in walking humans
2. a demonstration of the scientific potential of our approach in a study comparing the circadian rhythms of two bat species (*M. bechsteinii* and *N. leisleri*)
3. the development of a toolset (trained models, software, data, tutorials) for broader application in animal tracking research

### A random forest model to classify activity states based on automatically recorded VHF signals

From 2018 to 2021, we operated a network of 15 custom-designed automatic radio-tracking stations (henceforth ‘tRackIT stations’; (Gottwald et al., 2019; Höchst et al., 2021) in the Marburg Open Forest, Hesse, Germany (Fig. 2). This mixed temperate forest of 200 ha is dominated by European beech (*Fagus sylvatica*) and is home to 13 species of bats and 43 species of birds. Each tRackIT station consisted of four directional antennas, oriented north, east, south and west, and was powered by solar panels.

**Fig. 2.**
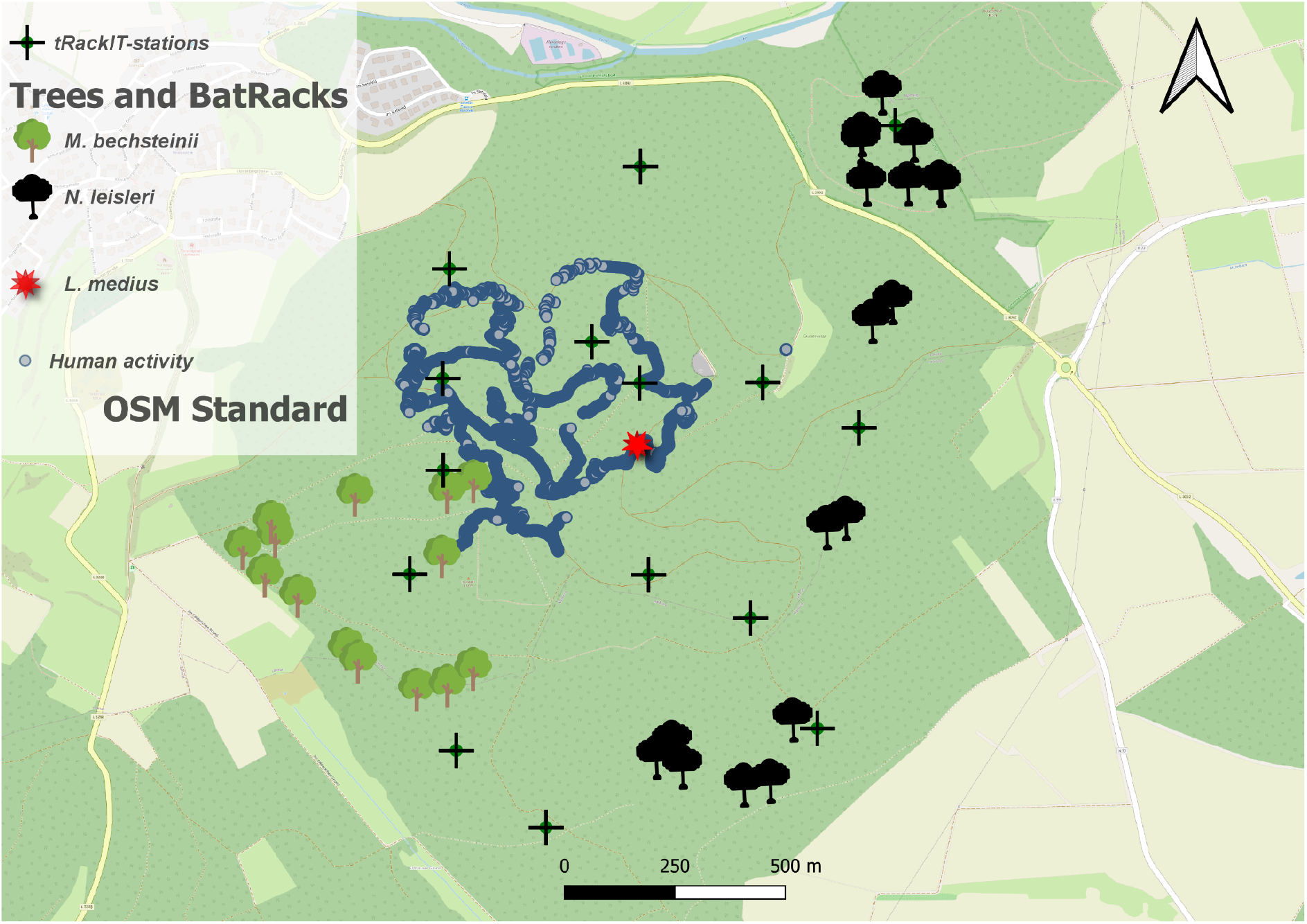
The Marburg Open Forest in Hesse, Germany. The map shows the locations of the tRackIT stations (Gottwald et al., 2019, Höchst & Gottwald et al., 2021), the roost trees of bats (*M. bechsteinii,N. leisleri)* observed by BatRack multi-sensor stations (Gottwald & Lampe et al., 2021), the breeding site of a woodpecker (*L. medius*) and the GPS track (shown in blue) of the activity simulation used to test the transferability of the classification method to birds and humans. (© Openstreet MAP OSM)

Every year, bats were caught and then tagged with customised VHF transmitters of different sizes and weights (V3+, Dessau Telemetrie-Service; see S2 for technical details, methods and permits). In total, 91 bat individuals from two focus species were captured and tagged (66 *M. bechsteinii* and 25 *N. leisleri*). Transmitter signal frequency, duration and strength as well as the timestamp of the signal of all individuals tagged at a given time were simultaneously and automatically recorded. Beginning in 2021, the data were transferred in real-time from the field to a streaming database hosted at Philipps-Universität in Marburg (Höchst & Gottwald et al., 2021; https://github.com/Nature40/tRackIT-OS). The patterns in the strength of the recorded signals were used together with ML algorithms to classify the activity of the tagged individuals.

Tags that provided inaccurate data, either because of a station configuration error or because too few signals were received, were excluded (n = 15, see the detailed flowchart in Fig. S3), from both model training and ecological study. For the two individuals retagged within the same year, the second monitoring period was excluded to ensure that all individuals were equally naïve to the tagging procedure. In total, data from 72 individuals (*M. bechsteinii*: N_ID_ = 52, N_Obs_ = 577 977; *N. leisleri*: N_ID_ =20, N_Obs_ = 204 443) monitored for 19 days on average (according to battery power) were used to distinguish active from passive states. In addition, 23 of the 72 individuals (6 *N. leisleri* and 17 *M. bechsteinii*) were monitored using a multi-sensor tool (Gottwald et al., 2021) to generate groundtruth for model training.

### Groundtruth

Supervised machine learning requires training and test data for implementation. To do so, we supplied the random forest model with periods of known activity and inactivity from tagged individuals of two bat species. Part of the labelled data were used to complete the training phase and the remainder held back for testing.

First, the roost trees of tagged bats were located via manual radio-telemetry between June 9, 2020 and July 26, 2020 and between May 10, 2021 and August 18, 2021. Custom-made video recorder (‘BatRack’) units were then set up to automatically record videos of tagged individuals (Gottwald et al., 2021); https://nature40.github.io/BatRack/ (vid 2)). BatRacks consist of a VHF antenna and an infrared video unit connected to a Raspberry Pi single board computer. The cameras were installed with a focus on the roost entrance and its surrounding area (40-m radius), which allowed the motion of tagged individuals to be captured on the video tracks. The infrared camera unit was automatically triggered by the VHF signal of the bat transmitters and started recording if the VHF signal strength exceeded a threshold of -60 dBW, i.e. when a tagged bat flew close to the roosting tree and the BatRack system.

The video tracks recorded by BatRack units were manually reviewed in conjunction with the VHF signal, and the observed behavioural sequence were then classified into the categories swarming, passing, entering or emerging from the roost. Sequences that showed swarming, passing or emerging were classified as active, and the time between entering and emerging from the roost as inactive. In addition to the sequences recorded on video, periods of time were classified as active if an individual was recorded in short time intervals on widely separated VHF receivers (tRackIT stations and BatRacks). From the three (2020) to nine (2021) BatRacks set in front of a total of 30 roosting trees of 6 *N. leisleri* and 17 *M. bechsteinii* individuals (Fig. 2), 723 h of behaviour were recorded. For these time periods of known activity type, a passive or active label was assigned to the VHF data recorded by one or more of the 15 radio-tracking stations.

### Predictor variables

We calculated 29 predictor variables thought to capture the patterns in the signal strengths over time by applying rolling windows of ±10 data entries, corresponding to an approximate time window of 20 s each, to the classified VHF data recorded by the tRackIT stations. To smooth out noise or potentially distracting fluctuations in the signal, a Hampel filter was calculated, in which data points that differ from the window median by more than three standard deviations are replaced by the median (Hampel, 1974). A mean and a max filter on the raw data of the main receiver was also applied. Next, the variance, standard deviation, kurtosis, skewness and sum of squares were calculated for both the raw and the smoothed data, to capture the variability and shape of the data distribution within the window.

Only one antenna is necessary to classify VHF signals into active vs. passive states (Kays et al. 2011). However, agreement between receivers of the same station provides additional information and can improve the reliability of the classification. This is especially likely if the individual is relatively close to the station (< 400 m in our scenario). When data from two receivers of the same station were available, the variance in the signal strength between receiver 1 and receiver 2 was calculated together with the correlation coefficient and the covariance of the signal strength in a rolling window of ±10 data entries. All variables are described in Supplement S1.

### Training and test data

The groundtruth dataset was balanced by randomly down-sampling the activity class with the most data to the amount of data contained by the class with the least data. These balanced datasets were then split into 50% training data and 50% test data for data originating from one receiver. The same procedure was used for data derived from the signals of two receivers, resulting in two training and two test datasets. From a total of 3,243,753 VHF signals, 124,898 signals were assigned to train the two-receiver model and 294,440 signals to train the one-receiver model (Table 1). The datasets are available at: https://doi.org/10.17192/fdr/80.

**Table 1.** Characteristics of the test and training data obtained from 723 h of video observation on 23 tagged individuals.

| Setup | Active data | Passive data | Total data points | Balanced active (train/test) | Balanced passive (train/test) |
| --- | --- | --- | --- | --- | --- |
| 1 receiver | 588,880 | 2,654,873 | 3,243,753 | 294,440 | 294,440 |
| 2 receivers | 249,796 | 1,469,674 | 1,719,470 | 124,898 | 124,898 |

### Model tuning

A random forest model was chosen as the classification method because it tends to perform better than other classifiers, as shown in an extensive comparative study (Fernández-Delgado et al., 2014). This model type is also robust against multicollinearity in predictor variables, especially when used with feature selection procedures (Gregorutti et al., 2017), as was the case in our approach. Since not all variables are equally important to the model and some may even be misleading, 50% of the data recorded by either one or two receivers were used to perform a forward feature selection as implemented in the “CAST” package (Meyer et al., 2018). This resulted in two random forest models, for data collected by one receiver and two receivers, respectively. The data and code are available at https://doi.org/10.17192/fdr/80.

### Test data for birds and humans

First, a series of 61 controlled walks was conducted with human volunteers to test the reliability of the trained models when applied to various activity patterns (resting, small-, medium- and large-scale movements) and tag positions. This was achieved by moving the VHF transmitters at two different heights, 15 cm above the ground at the ankle and 4 m above the ground, on a pole attached to a backpack, around the tRackIT stations. Resting was simulated by standing still. Movements on a small spatial scale were simulated by walking and hopping back and forth over an area of about 1 m^2^. Movements at a medium spatial scale were simulated by walking within areas of 40 m^2^, with multiple back and forth displacements and displacements of at least 200 m were used to simulate large-scale movements. Each movement type was performed for 3–10 min at different positions within the northwestern part of the study area (Fig. 2). The beginning and end times of the sequences were recorded and all signals simultaneously recorded by one or more of the 15 tRackIT stations, were then manually assigned to their known activity type (active or passive, depending on the type of movement simulated).

To test the reliability of the model on birds, a transmitter was attached to the back of a middle spotted woodpecker (*Leiopicus medius*) (see Supplement S2 for technical details and permits) and a daylight variant of the BatRack (“BirdRack”) was placed in front of its nesting tree for 4 consecutive days. A typical recorded sequence consisted of flying, hopping up the stem and a very short feeding sequence during which the bird remained motionless at the entrance of its breeding cave. Since the feeding sequence was usually shorter than three consecutive VHF signals (∼2.5 s), all recorded signals within such an sequence were classified as active. To generate sufficient inactive sequences, 2,200 random data points were sampled from signals recorded by tRackIT stations each night between 0:00 h and 2:00 h, while the woodpecker was asleep, over four consecutive nights. The human activity dataset consisted of 32,175 data points (26,133 active, 6,042 inactive) and the dataset of the woodpecker, based on the 75 observed activity sequences, of 17,541 data points (8,741 active, 8,800 inactive).

### Model validation

The trained random forest models were applied to the 50% of the data withheld for testing to evaluate their performance in classifying bat activity. The same trained models were applied to the bird and human activity datasets after the same predictor variables as used for the bats had been calculated. In a first step, the true positive rate (TPR) was calculated as the ratio of correctly identified incidents to all incidents, based on a comparison of the observed data with the activity class attributed by the random forest models for all dataset types (i.e. human activity dataset, woodpecker dataset and bats test-dataset). In a second step, the F-scores were calculated as the harmonic mean of the precision (*true positives / (true positives + false positives)*) and recall (*true positives / (true positives + false negatives)*). This value varies between 0 and 1, with values close to 1 indicating that the model has precision and recall values close to their maxima (Chinchor, 1992). Finally, the circadian activity patterns of the woodpecker were compared with the pattern expected for diurnal vertebrates.

The results of the ML-based approach were compared with those of a threshold-based approach by calculating the difference in the signal strength between successive signals for all three test datasets (bats, bird, humans). We applied a threshold of 4 dB which was deemed appropriate to optimally separate active and passive behaviours in previous studies (Holland et al., 2011). In addition, the optimize-function of the R-package stats (R Core Team, 2021) was used to identify the value of the signal strength difference that separated the training dataset into active and passive with the highest accuracy. This value was also applied to all three test datasets.

Both the data used in the validation and the code are available at: https://doi.org/10.17192/fdr/82.

## Results and discussion

### Random forest model to classify the activity of vertebrates of different sizes and movement types

The trained random forest models performed equally well, with F-scores of at least 0.96 and TPRs no less than 0.95, when applied to the validation data (Fig. 3 (a)). Whether the tag was positioned 15 cm or 4 m above the ground had no impact on the classification accuracy. The four activity levels simulated by humans were detected similarly well, with TPRs between

**Fig. 3.**
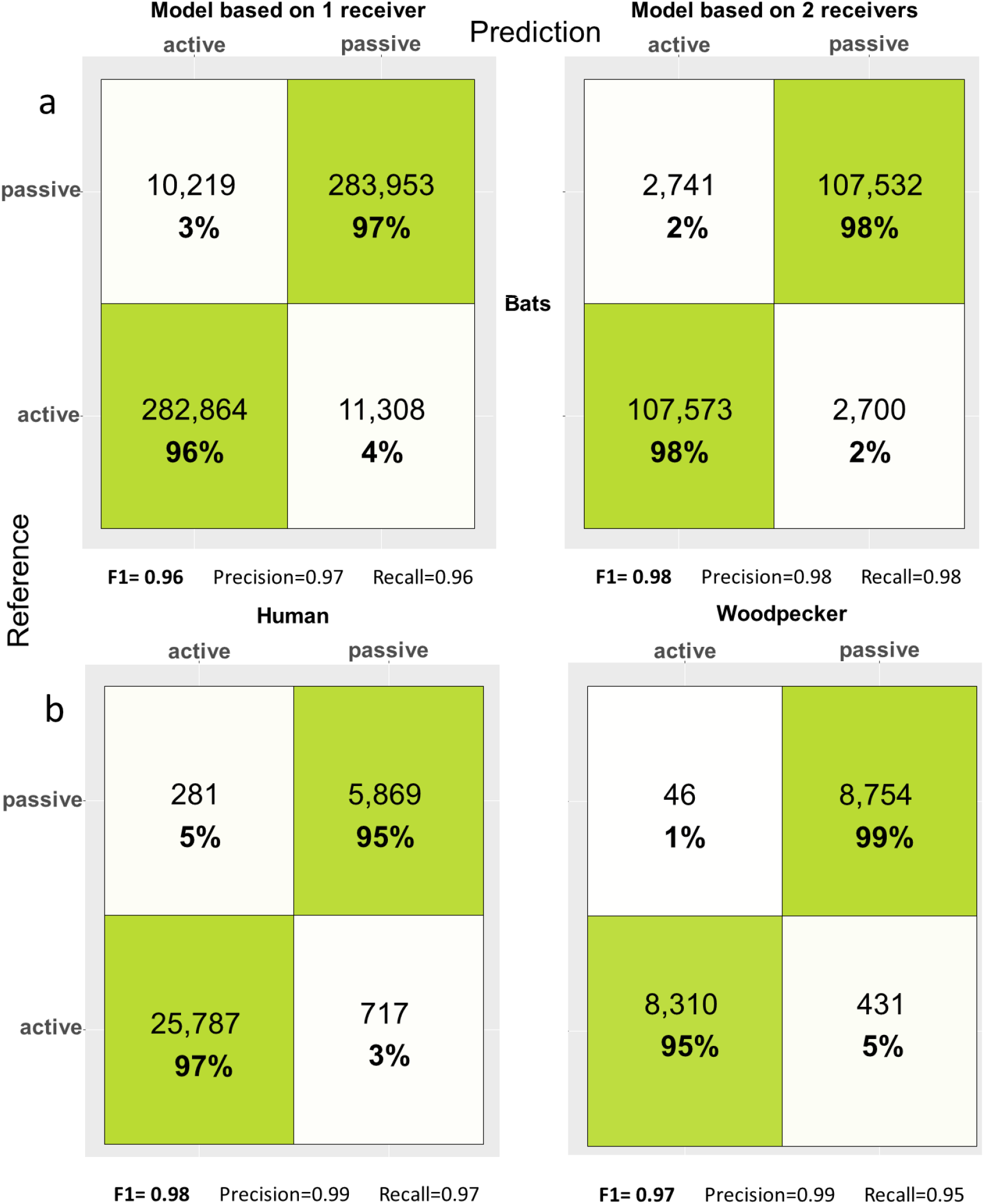
(a) Confusion matrix for the one-receiver- (left) and two-receiver (right) models. (b) Confusion matrix for different activity levels simulated by humans carrying transmitters (left) and based on the observed behaviour of a middle spotted woodpecker (right).

0.95 and 0.97. There were also no differences between the activity data of humans and those of the woodpecker (Fig. 3 (b)). Visual assessment of the active / passive sequences for the woodpecker showed typical patterns of high activity during the day, starting around sunrise (05:12) and ending around sunset (21:30; Fig. 4). For the results of the variable selection procedure, see Supplement S1.

**Fig. 4:**
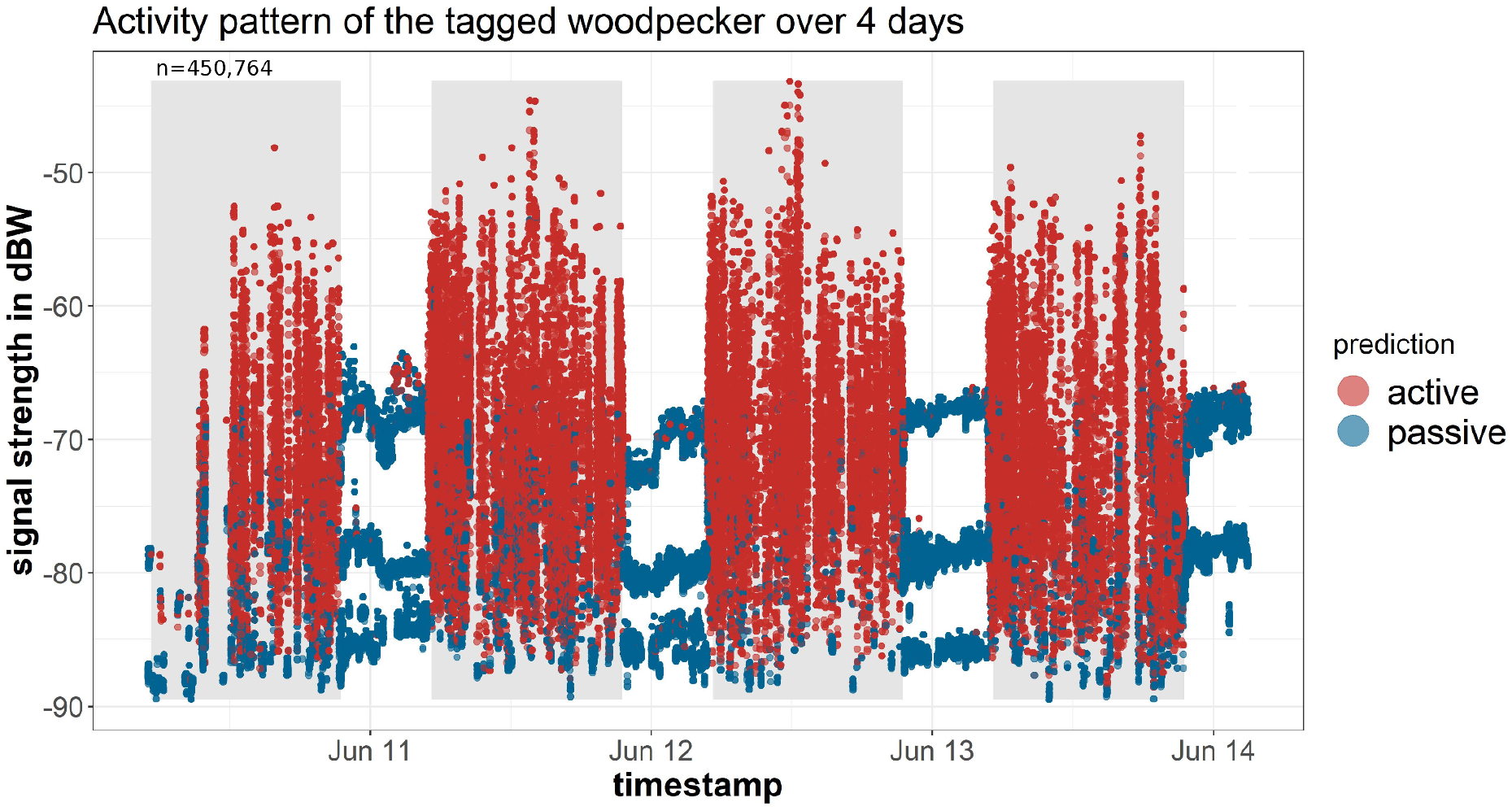
Signal strength [dbW] from a woodpecker tagged over four consecutive days and nights and the corresponding classification of the bird’s activity into active (N = 146,962) and passive states (N = 303, 802). Periods of high activity were consistent with the diurnal (5:00 to 21:00, grey background) activity patterns expected for this species.

By contrast, the threshold-based approach, using a 4-dB signal strength as the separation value as proposed in the literature (Holland et al., 2011), resulted in F-scores of 0.49, 0.68 and 0.7 for the test data of bats, woodpecker, and human activity respectively. A threshold of 1.3 dB, determined by optimising the separation of the training data, yielded F-scores of 0.72, 0.83 and 0.71 for bats, woodpecker and human activity respectively.

The results show that a threshold-based approach is generally able to classify active and passive behaviour. However, the proposed default threshold of 4 dB does not optimally separate active and passive behaviour and even a value calibrated using a groundtruth dataset suffered from a high variance in the performance metrics when applied to the different test datasets. In comparison, the random forest model classified the test data with high accuracy and low variability.

The trained models are available at: https://doi.org/10.17192/fdr/79.

### Ecological case study: comparison of activity patterns in two forest bat species

In the following, an ecological case study is presented to highlight the advantages of the fine-scale classification of activity states at a 1-min rate for two species monitored over four consecutive years. Both *M. bechsteinii* and *N. leisleri* are protected species (Habitats Directive 92/43/EEC) endemic to Eurasian forests but they differ substantially in their foraging habits. *N. leisleri* feeds on ephemeral insects that occur in large numbers, but only for short periods at dusk and dawn (Beck, 1995; Rydell et al., 1996) while *M. bechsteinii* partially collects its prey from the vegetation (Dietz & Pir, 2011; Kerth et al., 2001) and is thus generally less dependent on the timing of insect flight activity (Rydell et al., 1996).

The questions posed in this research were: 1) Do the two bat species differ in their overall probability of activity? 2) Do they differ in their timing of activity over the course of their circadian rhythms 3) How do activity patterns differ as a function of a species’ reproductive status (e.g. in lactating vs. breeding vs. non-reproducing individuals)? To answer these questions, we compared the timing of the onset and end of activity periods, the timing of maximum activity and the overall duration of night-time activity bouts using the data processed with the random forest model.

### Data collection and processing

The data were processed using the corresponding tRackIT R package (https://github.com/Nature40/tRackIT), which provides all functionalities needed for the classification of activity. The signals of individuals were filtered from the raw data based on frequency, signal length and the start and end of the tagging period. To apply the random forest models the predictor variables depending on the number of receivers were calculated. Each data point was then classified as active or passive behaviour using the appropriate model. All validation steps described in the section ‘Model validation’ were performed on individually classified data points. The classified data were then aggregated into 1-min intervals by extracting the most frequent class (active / passive) from all data points within each interval. The complete dataset and the code (284 GB) used for processing are available at: https://doi.org/10.17192/fdr/83. Exemplary data processing using the tRackIT package, shown with a small dataset, can be found at: https://doi.org/10.17192/fdr/81.

### Statistical analyses

All analyses were conducted with R v. 4.1.2 (R Core Team, 2021), using the mgcv package for additive models (Wood, 2011). Reproducible scripts are available at: doi/dhtfjzgi.

Hierarchical generalised additive models (HGAM) were used to compare differences in the overnight activity patterns of *M. bechsteinii* and *N. leisleri*. These classes of models can be applied to estimate non-linear relations between responses while allowing for a variety of error terms and random effect specifications (Bogdanović et al., 2021; Pedersen et al., 2019). In this study, activity was modelled over the course of the 24-h cycle as shown in Eq. 1:

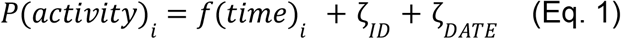

where the probability of activity for observation *i* is modelled as a binomial variable (0: inactive, 1: active) as a function of the time of day, which. was centred around sunset to account for seasonal shifts in daylight. A circular cubic spline was specified to constrain the beginning and end of the 24-h cycle so that they matched. Individual identity and date were added as random effects to account for individual, seasonal and yearly effects. Given the volume of data (> 700,000 observations), all models were fitted through the bam() function for faster model estimation.

Given the short timespan between observations, it was necessary to account for autocorrelation. This can be done with the bam() function but it has to be defined manually. Thus, in a first step the model was fitted without an autocorrelation structure and the start_value_rho() from the itsadug package (van Rij et al., 2020) was used to estimate autocorrelation (**ρ**) among residuals at the first lag (**ρ** = 0.57). The model was then refitted with the estimated autocorrelation value with an AR1 structure. This procedure successfully accounted for autocorrelation, as evidenced by the decrease in the median autocorrelation to −0.13 in the refitted model. Visual inspection of the autocorrelation confirmed that rho remained < |0.15| at all lags.

Differences in the activity patterns of the two species were evaluated by comparing the Akaike information criterion (AIC) values between a model in which species did not vary in their daily activity patterns (Model 0) against one in which the effect of time of day varied between species (Model 1, using the “by = species” argument to specify a time × species interaction). Activity parameters among species were also compared by calculating the difference in spline functions, Δ *f(time)*, as the difference between *M. bechsteinii* and *N. leisleri* splines. This more precisely revealed the period of the day when the two species were most likely to differ in their probability of activity (negative value: P(activity)_Bechstein_ < P(activity)_Leisler_, positive value: P(activity)_Bechstein_ > P(activity)_Leisler_). Because *N. leisleri* could not be monitored during the 2020 field season, due to the Covid-19 pandemic, the data were also analysed by removing the 2020 data. Since there was no difference in the inference or support for the best model, the results from the full dataset are presented (see the Supplement for the complete analysis).

The activity patterns of the two bat species were further characterised by calculating the following metrics based on the predicted values for Model 1:

- Onset and end of activity periods, defined as the first and last time of day when the probability of activity was larger than chance (i.e. p(activity) > 0.5)
- Time of peak activity, calculated as the time of the day when the probability of activity was maximal
- Activity density, defined as the area under the curve between the onset and end of activity

A similar approach was used to compare the effect of reproductive status on the activity patterns within species. This was done using females, as their reproductive status at capture was consistently monitored over the 4 years of sampling. The transition between reproductive periods was determined based on the reproductive characteristics observed during capture events, carried out two to three times a week. The lactation period was defined as beginning with the capture of the first lactating female of the respective species and the post-lactation period as coinciding with the capture of the first fledged juvenile (*M. bechsteinii*: N_ID_ = 35, N_Obs_ = 384 536; *N. leisleri*: N_ID_ = 19, N_Obs_ = 203 261). Separate models were then run for each species and models with and without the inclusion of reproductive status, as an interaction term with time of day, were compared. It was expected that reproductive individuals (pregnant or lactating) would have longer activity periods overall. Lactating individuals were also expected to have more variable patterns of nocturnal activity as they need to frequently interrupt their activity to feed their pups at the roost site (Dietz & Kalko, 2007; Lučan & Radil, 2010).

## Results and discussion

### Species comparisons of circadian activity

*Nyctalus leisleri* and *M. bechsteinii* showed pronounced differences in the shapes of their activity curves, as indicated by AIC model selection (ΔAIC = 15092, Table 2; Fig. 5). While both species appeared to synchronise their onset of activity with sunset, *N. leisleri* was active an average of 19 min earlier than *M. bechsteinii. N. leisleri* also reached peak activity earlier, but its activity markedly declined as soon as *M. bechsteinii* became highly active. The latter species was highly active throughout most of the night, as indicated by a significantly higher activity density (area under the curve when p(activity) > 0.5 [95% CI]; *M. bechsteinii*: 4.70 [4.56; 4.83]; *N. leisleri*: 3.42 [3.23; 3.62]). However, *M. bechsteinii* reached the end of its activity period an average of 12 min sooner than *N. leisleri* (Table 3).

**Table 2.** Model coefficients (β) and standard errors (SE) and test statistics (z, p) for the linear portion of the additive models (intercept) along with smoothed parameters for nonlinear terms (edf: estimated degrees of freedom, chi-squared and p-values).

| | Model 0 (AIC = 263401.1; $R^2$ = 0.45) | | | | | | Model 1 (AIC = 248309.1; $R^2$ = 0.47) | | | | |
| --- | --- | --- | --- | --- | --- | --- | --- | --- | --- | --- | --- |
| Linear terms | $\beta$ | SE | z | p | | Linear terms | $\beta$ | SE | z | p | |
| Intercept | -1.74 | 0.13 | -12.98 | $< 2 \times 10^{16}$ | | Intercept | -1.74 | 0.13 | -13.29 | $< 2 \times 10^{16}$ | |
| Smoothed terms | edf | df | $\chi^2$ | p | % Variance | Smoothed terms | edf | df | $\chi^2$ | p | % Variance |
| time | 51.84 | 118 | 4061287 | $< 2 \times 10^{16}$ | 17.42 | time:Leisler | 45.82 | 118 | 188033 | $< 2 \times 10^{16}$ | 8.83 |
| | | | | | | time:Bechstein | 47.23 | 118 | 3571852 | $< 2 \times 10^{16}$ | 22.46 |
| Random effects |  |  |  |  |  | Random effects |  |  |  |  |  |
| ID | 67.07 | 71 | 1976322 | $< 2 \times 10^{16}$ | 10.18 | ID | 66.32 | 71 | 1824189 | $< 2 \times 10^{16}$ | 4.78 |
| DATE | 280.26 | 306 | 3084452 | $< 2 \times 10^{16}$ | 17.44 | DATE | 282.01 | 306 | 2421809 | $< 2 \times 10^{16}$ | 10.73 |

**Fig. 5.**
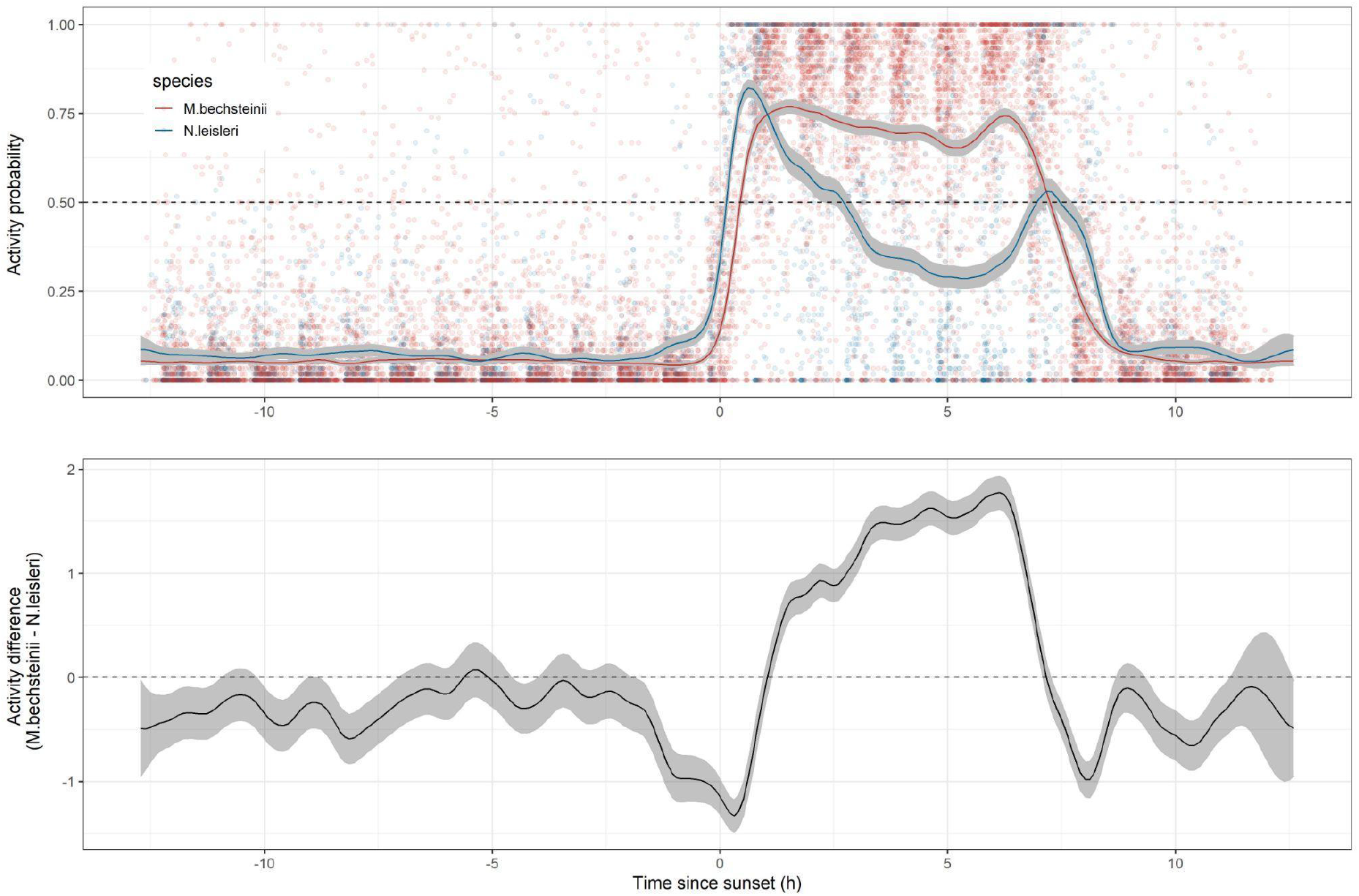
*Nyctalus leisleri* was consistently active sooner than *M. bechsteinii*, but the latter species had longer periods of continuous activity. Top panel: the points represent the activity probability calculated over 1-h intervals, and the solid lines the predicted values from the best HGAM model. The dashed line indicates the times when the population was equally likely to be detected as active or passive. Bottom panel: difference in the activity probability calculated from the best HGAM model. Positive values indicate a larger activity probability for *M. bechsteinii* than for *N. leisleri*.

**Fig. 6.**
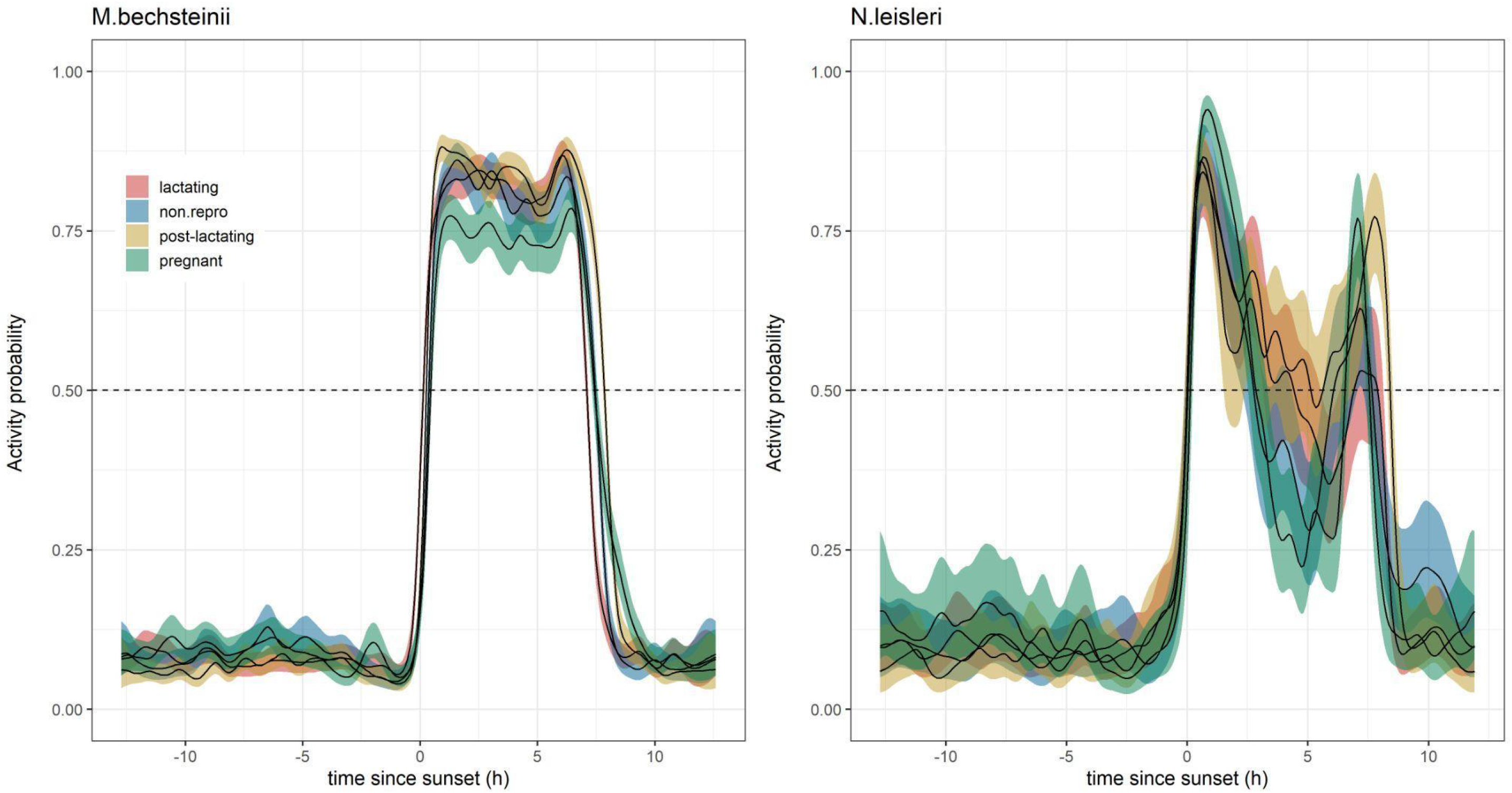
Pregnant females of *M. bechsteinii* had a lower activity probability while for *N. leisleri* post-lactation resulted in extended activity bouts around sunrise. Solid lines represent the predicted values of the HGAM models fitted separately to each species. The dashed lines indicate the times when the population was equally likely to be detected as active or passive.

**Table 3.** Activity metrics of *N. leisleri* and *M. bechsteinii*. Wake-up and sleep times were calculated as the first and last time of day when the probability of activity was > 0.5. The time of peak activity represents the time of day when the probability of activity was maximal. The activity density was calculated as the area under the curve between wake-up and sleep times.

| Metric | *N. leisleri* | *M. bechsteinii* |
| --- | --- | --- |
| Activity onset (h) | 00:12 | 00:31 |
| Time of peak activity (h) | 00:37 | 01:33 |
| Activity end (h) | 07:23 | 07:11 |
| Peak P(activity) | 0.82 [0.80; 0.84] | 0.70 [0.75; 0.79] |
| Activity density | 3.42 [3.23; 3.62] | 4.70 [4.56; 4.83] |

These results are generally in line with previous observations of the activity patterns of *N. leisleri* (Ruczyński et al 2017, Shiel et al 2007). No comparable studies exist for *M. bechsteinii*, but in acoustic studies with results reported at the genus level all-night activity was determined for *Myotis* (Perks & Goodenough, 2020). However, our study is the first to investigate the overlap of these two species within the same study area. Specifically, we were able to show distinct activity patterns for these two species, characterised by a slight shift in their timing of activity. Given that these species have evolved to occupy different ecological niches, these patterns are much more likely due to a synchronisation of activity peaks with prey abundance rather than to an avoidance of competition (Ruczyński et al., 2017). *N. leisleri*, like other aerial hawking bats, has likely evolved to exploit insect emergence at dusk and dawn, thus avoiding the greater predation risk that may occur at higher light levels (Rydell et al., 1996). By contrast, *M. bechsteinii* and other gleaning bats are less constrained to flying insects as a food source such that an onset of activity comparable to that of *N. leisleri* would not bring substantial additional benefit.

### Variation in circadian activity according to the breeding status of individuals

Reproductive status had a significant influence on the activity probability (*M. bechsteinii*: ΔAIC = 3657; *N. leisleri*: ΔAIC = 2974), with marked differences between the two bat species (Fig. 5). In *M. bechsteinii*, lactating and post-lactating females behaved very similarly to non-reproductive females while pregnant females had a lower activity probability during the night. All individuals were highly synchronised with respect to activity onset, regardless of reproductive status, but post-lactating females had longer activity periods than the other groups of females. In *N. leisleri*, the major differences were between pregnant and post-lactating females. Thus, pregnant females concentrated their activity within 1 h after sunset and decreased their activity more rapidly than post-lactating females.

Although an effect of reproduction on the probability of activity was supported, it was not as expected. For both species, reproductive individuals seemed to end their activity period quicker than post-lactating and non-reproductive females. In addition, while pregnant females had lower activity levels, lactating females had neither more variable patterns of activity nor longer activity periods, contrary to what we expected. Note that pregnancy does not always result in lower activity levels, as shown in *Myotis daubentoni* (Dietz et al. 2007). By contrast, females of both *Nyctalus noctula* and *N. leisleri* had significantly shorter activity times during pregnancy than during the lactation period (Ruczyński et al., 2017).

Our results demonstrate that significant differences in the timing of activity according to a species’ ecological preferences or seasonality can be detected using the system presented herein.

## Conclusion

Using a large dataset consisting of the observed behaviours of tagged bat individuals, we trained two random forest models to classify novel data from the same species into fundamental behaviours, and with high precision and high temporal resolution (∼1 sec interval). The amount of data used to train the models was large enough to sufficiently represent the diversity of potential signal paths and patterns, thus ensuring the applicability of the models to taxa with different movement behaviours. This was demonstrated by the comparable precision achieved when the same models were applied to groundtruth data from another flying species (woodpecker) and to a walking vertebrate (humans). This strongly suggests that our method generalises well and can be applied to a wide variety of small to medium-sized vertebrates with similar accuracy. By contrast, the precision of the threshold-based approach was consistently lower and the variability between the different test datasets higher.

Whether different behaviours can be recognised, as is possible with accelerometers, remains to be determined. The fact that the variance in the signal pattern depends less on the intensity of the movement than on the signal path poses a problem. The spatial context of the receiving stations as well as the localisation algorithms presented in Gottwald et al. 2019 could provide additional information.

A comparison of the activity probabilities of two ecologically different bat species showed that our method can detect subtle differences not only between species (differences in activity onset of < 20 min) but also between different reproductive states within a species. A more in-depth analysis of activity bouts as a function of abiotic factors or the detection of changes in patterns indicating, for example, the transition from non-breeding to breeding, has not been conducted here, but such studies are likely to be feasible.

The tRackIT system can currently record up to 90 individuals at a time within the same spatial context, but technology that allows for higher numbers is under development. Given the relatively low costs of the transmitters (∼130 €) and tRackIT stations (∼1500 €), the monitoring of an entire community of small forest vertebrates at high temporal resolution becomes possible. The high precision and high temporal resolution of our approach may also open new research avenues on the variations in the activity patterns among and within species that reflect their response to the environment. The tRackIT system is currently being used for nature conservation as a means to narrow down the time of death of chicks of meadow-breeding birds (https://www.audi-umweltstiftung.de/umweltstiftung/de/projects/greenovation/telemetry-technology.html).

With the recent advances in open-source automatic radio-tracking (Gottwald et al., 2019; Höchst et al., 2021) together with the data-processing functionalities of the tRackIT R-package, the scientific community is now equipped with an accessible toolset that allows the activity patterns of small animals to be analysed and classified at high temporal resolution. The scientific insights that can be expected from such studies have the potential to deepen our understanding of the ecology and behavior of small animal species in unprecedented ways (Nathan et al., 2022).

## Author contributions

JG planned the study and organised the field work for collecting groundtruth and bat activity data. JG, LL, JM, TG (bats and humans), DGS, SR, KL, MB (woodpecker), PL, JG and JH (hardware and software development and implementation) participated in the fieldwork effort required for locating, tagging and monitoring the bat and bird populations. BF, RB, TM, NF and TN supervised the study. JG developed the associated R-Package and defined the research questions in association with DGS, SR and RR. JG applied and then validated the machine learning model. RR was responsible for the statistical analysis of the activity patterns of bats. JG and RR jointly wrote the first draft of the manuscript. All authors contributed to subsequent versions.

## Acknowledgements

The research was funded by the Hessen State Ministry for Higher Education, Research and the Arts, Germany, as part of the LOEWE priority project Nature 4.0 – Sensing Biodiversity (https://uni-marburg.de/natur40). We thank all Master and Bachelor students who supported the fieldwork. Special thanks to C. Kodges and F. Strehmann for their tireless work in catching and tagging the birds and bats.

## Data and code availability

To ensure the complete reproducibility of our research, all data and the code used are stored in a data collection at data_UMR, the research data repository of Philipps-Universität Marburg (https://data.uni-marburg.de/; search: “classification of activity states in small vertebrates”).

In addition, to equip the research community with the novel tool introduced in our study, we developed two detailed tutorials, providing (1) all necessary steps required to translate the recorded raw VHF signals to an active/passive classification and (2) all analytical steps of the ecological case study. As the total dataset is > 287 GB, we recommend downloading only the necessary partial datasets. In the following, we specify which dataset is necessary for which analysis.

### Trained models

For all working steps involving an actual classification of VHF signals into passive/active, the trained models should be downloaded and stored into the “extdata” folder of the installed tRackIT R-Package. The models can be downloaded at: https://doi.org/10.17192/fdr/79

### Model tuning

To understand the machine learning process that led to the trained models, we offer a download of the corresponding training data and the R code, available at: https://doi.org/10.17192/fdr/80

Since we used an elaborate feature selection procedure, the training process can take several days.

### Model validation

The models were trained on a very large dataset consisting of the recorded behaviour of tagged bat individuals. The model was tested using 50% of this dataset. In addition, transferability was tested on two completely independent datasets, one consisting of observations of a tagged woodpecker and the other of simulations of the activity levels of humans carrying transmitters. All three datasets, including the R-code to validate the method, can be found at: https://doi.org/10.17192/fdr/82

### Ecological case study

The complete dataset, including the raw data used in the ecological case study, as well as the R scripts can be found at: https://doi.org/10.17192/fdr/83

The dataset is 287 GB in size. Already-compiled datasets necessary for the reproduction of our analyses are in the repository for the tutorials (next section).

### Tutorials

Two tutorials are provided. The first shows which sequence of tRackIT R-Package functionalities finally leads to a classification of active/passive behaviour (tRackIT-Tutorial-for-activity-classification). The second ensures the reproduction of all statistical analysis steps from the ecological case study (bat_data_HGAM_tutorial). Data, html and rmd files can be found at: https://doi.org/10.17192/fdr/81

## Supplementary Materials

### S1: Variable selection

The three most informative variables for the classification of active and passive behaviour based on signals recorded by only one receiver were the standard deviation of the signal strength after data smoothing through a Hampel filter, the sum of squares of the max filtered signal strength and the max filtered signal strength. For data recorded by two receivers, the standard deviation of the signal strength difference of receiver one and receiver two was the most important, directly followed by the variance of the Hampel-filtered signal strength of the first receiver. The other selected variables were of marginal importance. The importance of each variable for the two models is reported in Supplement S1.

**Table S1:** Calculated variables and their importance for the activity classification

| Variable Name | Description | var.imp. 1<br>receiver | var.imp 2<br>receivers |
| --- | --- | --- | --- |
| max_signal | raw signal strength | 0 | 0 |
| max | max filtered signal strength of receiver 1 | 13.45 | 8.9 |
| mean | mean filtered signal strength of receiver 1 | 0 | 1 |
| hample | Hampel filtered signal strength of receiver 1 | 0 | 0 |
| cor_max | correlation of signal strength on 2 receivers | 0 | 0 |
| cov_max | covariance of signal strength on 2 receivers | 0 | 0 |
| diff | difference of signal strength of 2 receivers | 0 | 0 |
| diff_std | standard deviation of signal strength difference (diff) | 0 | 100 |
| diff_var | variance of signal strength difference (diff) | 0 | 0 |
| kurt | kurtosis of signal strength of receiver 1 | 0 | 0 |
| kurt_hampel | kurtosis of Hampel filtered signal strength of receiver 1 | 0 | 0 |
| kurt_max | kurtosis of max filtered signal strength of receiver 1 | 0 | 0 |
| kurt_mean | kurtosis of mean filtered signal strength of receiver 1 | 0 | 0 |
| skew | skewness of signal strength of receiver 1 | 0 | 0 |
| skew_hampel | skewness of Hampel filtered signal strength of receiver 1 | 0 | 0 |
| skew_max | skewness of max filtered signal strength of receiver 1 | 0 | 0 |
| skew_mean | skewness of mean filtered signal strength of receiver 1 | 0 | 2,3 |
| std | standard deviation of signal strength of receiver 1 | 0 | 0 |
| std_hampel | standard deviation of hampel filtered signal strength of receiver 1 | 100 | 0 |
| std_max | standard deviation of max filtered signal strength of receiver 1 | 0 | 0 |
| std_mean | standard deviation of mean filtered signal strength of receiver 1 | 0 | 0 |
| sumsq | sum of squares of signal strength of receiver 1 | 0 | 0 |
| sumsq_hampel | sum of squares of Hampel filtered signal strength of receiver 1 | 5 | 0 |
| sumsq_max | sum of squares of max filtered signal strength of receiver 1 | 20,3 | 13,6 |
| sumsq_mean | sum of squares of mean filtered signal strength of receiver 1 | 8 | 0 |
| var | variance of signal strength of receiver 1 | 0 | 0 |
| var_hampel | variance of Hampel filtered signal strength of receiver 1 | 0 | 97 |
| var_max | variance of max filtered signal strength of receiver 1 | 0 | 0 |
| var_mean | Variance of mean filtered signal strength of receiver 1 | 0 | 0 |

### S2: Capture methods and permits

Bats were captured with mist nets and radio-tagged with temperature sensitive VHF transmitters (V3, Telemetrie-Service Dessau, 0.35 g) between April and August each year. The weight of the tags was < 5% of every bat’s body weight, to minimise the effect on manoeuvrability, foraging effort and energetic costs (Aldridge & Brigham, 1988).

For the handling and tagging of the bats, a license was issued by the Nature Conservancy Department of Central Hessen (‘Obere Naturschutzbehörde Mittelhessen, Regierungspräsidium Gießen’, v54-19c 2015 h01).

Woodpeckers were caught using three mist nets (Ecotone Mist Net 716/12) set up in sequence. The transmitters had a weight ≤ 3% of the bird’s body weight (Caccamise & Hedin, 1985) and were attached on the bird’s back with a Rappole-Tipton-type harness (0.5-mm elastic cord) (Rappole & Tipton, 1990). The license to tag birds was granted by the Nature Conservancy Department of Central Hessen (‘Obere Naturschutzbehörde Mittelhessen, Regierungspräsidium Gießen’, v54-19c 2015 h01 MR 20/15 Nr. G 10/2019).

### S3: Workflow tag verification

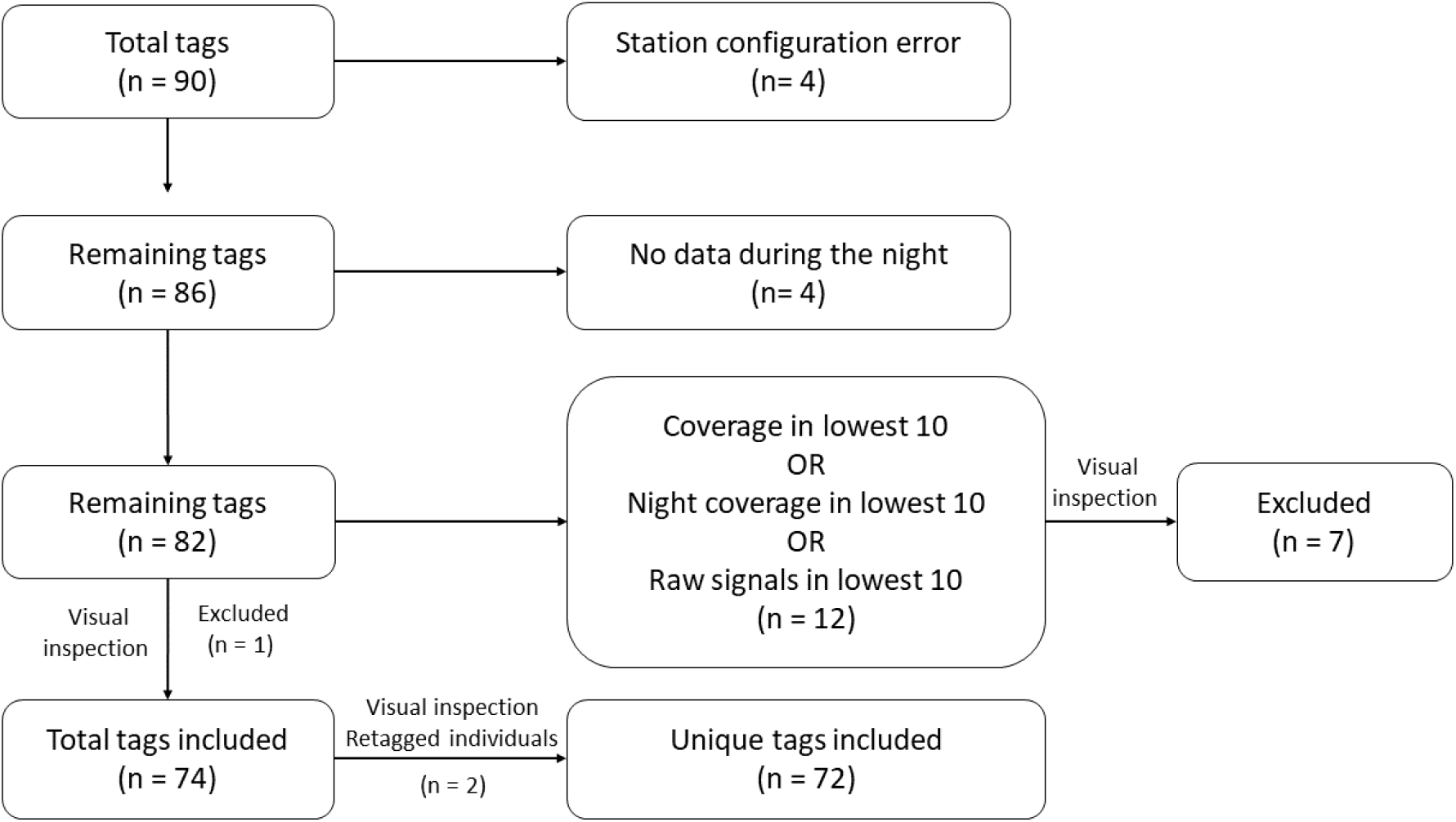

## Notes

https://github.com/Nature40/tRackIT-OS

https://nature40.github.io/BatRack/

https://doi.org/10.17192/fdr/79

https://doi.org/10.17192/fdr/80

https://doi.org/10.17192/fdr/81

https://doi.org/10.17192/fdr/82

https://doi.org/10.17192/fdr/83

### Competing Interest Statement

The authors have declared no competing interest.

